# Reducing the foreign body response on human cochlear implants and their materials *in vivo* with photografted zwitterionic hydrogel coatings

**DOI:** 10.1101/2022.11.28.518125

**Authors:** Ryan Horne, Nir Ben-Shlomo, Megan Jensen, Morgan Ellerman, Caleb Escudero, Rong Hua, Douglas Bennion, C Allan Guymon, Marlan R. Hansen

## Abstract

The foreign body response to implanted materials often complicates the functionality of sensitive biomedical devices. For cochlear implants, this response can reduce device performance, battery life and preservation of residual acoustic hearing. As a permanent and passive solution to the foreign body response, this work investigates ultra-low-fouling poly(carboxybetaine methacrylate) (pCBMA) thin film hydrogels that are simultaneously photo-grafted and photo-polymerized onto polydimethylsiloxane (PDMS). The cellular anti-fouling properties of these coatings are robustly maintained even after six-months subcutaneous incubation and over a broad range of cross-linker compositions. On pCBMA-coated PDMS sheets implanted subcutaneously, capsule thickness and inflammation are reduced significantly in comparison to uncoated PDMS or coatings of polymerized poly(ethylene glycol dimethacrylate) (pPEGDMA) or poly(hydroxyethyl methacrylate) (pHEMA). Further, capsule thickness is reduced over a wide range of pCBMA cross-linker compositions. On cochlear implant electrode arrays implanted subcutaneously for one year, the coating bridges over the exposed platinum electrodes and dramatically reduces the capsule thickness over the entire implant. Coated cochlear implant electrode arrays could therefore lead to persistent improved performance and reduced risk of residual hearing loss. More generally, the *in vivo* anti-fibrotic properties of pCBMA coatings also demonstrate potential to mitigate the fibrotic response on a variety of sensing/stimulating implants.

**Graphical Abstract:** 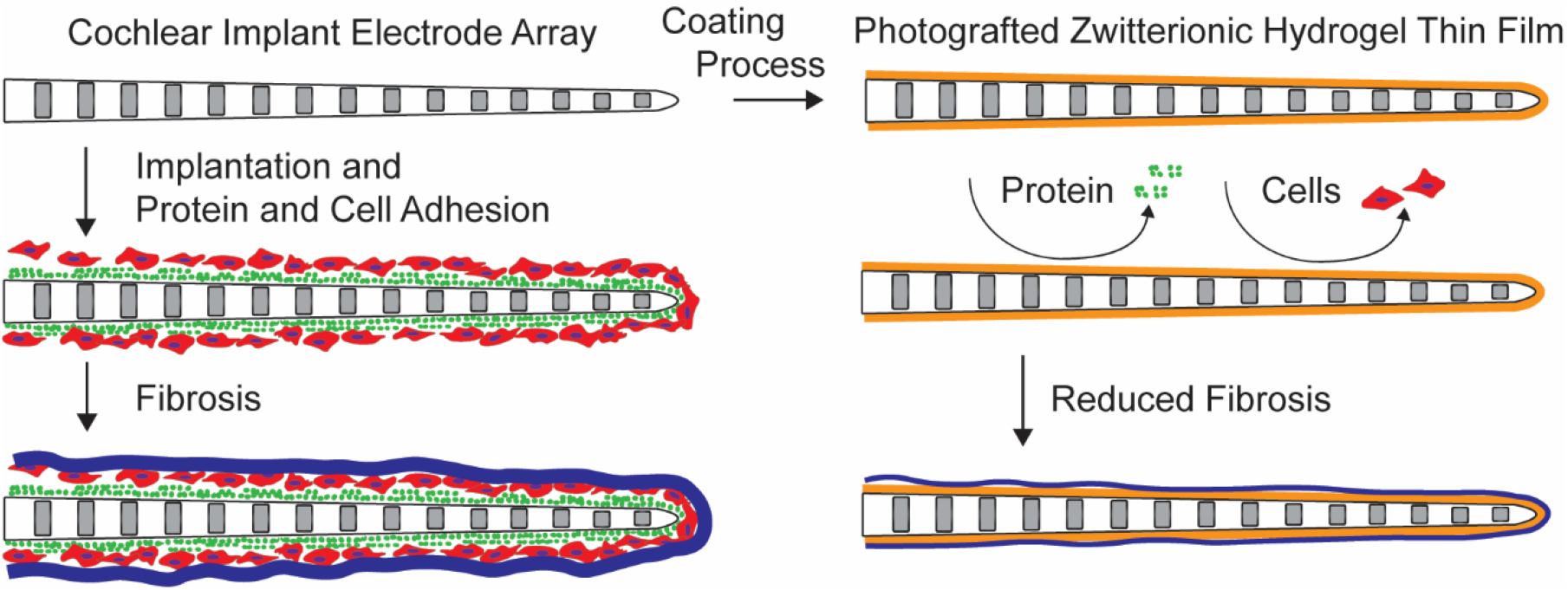

## Introduction

Implantable devices, which have wide ranging medical applications including cardiac pacemakers, deep brain stimulators, glucose monitors, and cochlear implants, have transformed patient health and outcomes. The utility of many of these biomedical devices, however, is limited by the biological responses to the foreign implant materials.[1–4] This foreign body response is characterized by persistent inflammation, macrophage infiltration and transformation to foreign body giant cells, and fibrotic capsule formation.[5] This immune response protects the *in vivo* environment from toxins, infection, chemical imbalance, cell lysis, and mechanical damage.[6] However, the foreign body response can also significantly impair the function of medically implanted devices. The inflammatory response can damage sensitive tissue structures, and fibrotic capsule formation can interrupt the intended interaction of the device with the host tissue. Encased in fibrotic scar tissue, these devices may not be able to stimulate, medicate, or sense with the desired degree of accuracy or precision.

For cochlear implants, if the stimulating electrode array becomes encased in fibrotic tissue through the foreign body response, the current flow is then spread, stimulating a wider population of target neurons, thereby resulting in decreased tonal specificity within the cochlea and diminished hearing outcomes.[7] Recently, hybrid cochlear implants have become critical for treating high frequency hearing loss in patients with preserved hearing at low frequencies.[8] However, the foreign body response in these patients can damage residual low-frequency hearing in addition to limiting the functionality of the cochlear implant to restore high-frequency hearing.[8, 9] In severe cases, the foreign body response can even lead to device malfunction or neo-ossification in the cochlea, preventing the possibility of reimplantation.[10]

The foreign body response evolves from initial protein adsorption to major tissue remodeling over the course of weeks to months. Immediately upon implantation, a foreign surface typically becomes deposited with ubiquitous *in vivo* proteins like fibrinogen and albumin.[11] Local immune cells, like neutrophils and later monocytes, recognize the deposited proteins and release pro-inflammatory signals.[12] As the response develops, macrophages adhere to the surface and attempt to engulf the foreign material, then gradually fuse into giant cells to form an immediate barrier around the implant.[12] Finally, fibroblasts are recruited and encase the giant cell layer in fibrotic tissue to further insulate the foreign material.[12]

If this cascade of protein deposition, cell adhesion and immune activation could be reduced, then the resulting giant cell and fibrotic capsule could be correspondingly decreased. Among the many approaches used to reduce the foreign body response, some have explored films of zwitterionic polymers or other hydrogel materials or to reduce protein deposition, cellular adhesion, and/or fibrotic capsule formation.[13–15] Zwitterionic polymers, consisting of repeating units that each carry paired positive and negative charges that balance to a net zero charge, are of particular interest as they create an internal dipole that strongly attracts water molecules in a highly ordered scaffold around the polymer.[16] This scaffold limits interaction with other species, thereby reducing protein and cell attachment to the surface.

While many of these zwitterionic polymer systems can be effective at mitigating the foreign body response, many lack the robustness required to comprehensively coat medical implants for long-term use. The modulus of hydrogels is often less than that of biological soft tissue, leaving the hydrogel vulnerable to damage as a stand-alone material. This effect can be mitigated by using the hydrogel as a coating, as long as it remains durable through handling, implantation and long-term use. Additionally, using zwitterions to permanently coat chemically inert surfaces, such as those used in cochlear implant electrodes can be difficult, because effective coating methods typically rely on interactions with or reactivity of the surface. Unfortunately, fibrosis on the cochlear implant or other neural interface electrodes limits device function by significantly reducing electrical signal transduction to sensory cells. Additionally, recent work has shown that the cochlear implant electrodes themselves are even more potent aggravators of the foreign body response than other materials such as the poly(dimethylsiloxane) (PDMS) housing.[17] One strategy has been to develop hydrogel coatings for cochlear implant materials that reduce protein and cellular adhesion, thus decreasing the capsule thickness in model systems.[18–20]

Our work has focused on simultaneously photopolymerizing and photografting cross-linked zwitterionic monomers, specifically carboxybetaine methacrylate (CBMA), to PDMS.[21] The resulting polyCBMA (pCBMA) forms a durable thin film that can coat a variety of surface types and demonstrates robust anti-fouling properties *in vitro* and against bacteria *in vivo*.[21–24] This strategy allows for dense packing of zwitterionic chains, creating a uniform hydrogel thin film firmly grafted to a surface. While this prior work shows promise, *in vivo* assessment is necessary to ascertain the true effect of photografted zwitterionic thin films on the foreign body response, as promising *in vitro* results are often not reproducible *in vivo*. Similarly, the effect of such a coating is best assessed on real medical devices, because different geometries, scales, materials, and handling impact the practicality of any surface modification.

The following studies assess the hypothesis that zwitterionic pCBMA hydrogel coatings reduce the foreign body response *in vivo* over both short and extended timeframes with specific focus on their application for cochlear implants. The longevity of the pCBMA coating is examined *in vivo* by imaging and quantitative maintenance of anti-fouling properties in a simple model system of PDMS sheets. The impact of hydrogel chemistry on *in vivo* fibrotic capsule thickness and the practicality of coated cylindrical implant geometry *in vivo* are assessed. Lastly, pCBMA thin films are applied to human cochlear implant electrode arrays subcutaneously *in vivo* to determine their capacity to reduce fibrotic capsules for one-year chronic implantations. Human cochlear implant electrode arrays are implanted subcutaneously in mice to enable the use of a small animal model to assess the effectiveness of an actual human device without the confounding effect of a traumatic insertion of a large device into the small, delicate mouse cochlea. These studies are designed to demonstrate the effectiveness of the coatings *in vivo* on commercial medical implants with multi-feature surfaces such as cochlear implant electrode arrays composed of platinum-iridium electrodes housed in PDMS.

## Materials and methods

### 2.1 Materials

Acetone and Irgacure 2959 (2-hydroxy-1-[4-(2-hydroxyethoxy) phenyl]-2-methyl-1- propanone were obtained from Sigma-Aldrich (St. Louis, MO). 3-([2-(Methacryloyloxy)ethyl]- dimethylammonio)propionate (CBMA) and PEG (molecular weight of 400, approximately 9 repeat units) dimethacrylate (PEGDMA) were purchased from TCI Chemicals (Portland, OR) and Polysciences (Warrington, PA), respectively. Different poly(dimethyl siloxane) (PDMS) systems used included silastic tubing (Dow Corning, Midland, MI) and non-reinforced medical grade silicone sheeting (0.01 in. thickness, Bentec Medical, Inc., Woodland, CA). Benzophenone was purchased from Acros Organics (NJ). An Omnicure S1500 lamp (Excelitas Technologies, Mississauga, Canada) was used for photocuring under pure nitrogen gas (Linde, Inc., Danbury, CT). The surfactant (Gelest, Morrisville, PA) used was dimethylsiloxane-acetoxy terminated ethylene oxide block copolymer (75% non-siloxane) and acted as a dispersing agent for coating cylindrical geometries.

### 2.2 Zwitterionic thin film fabrication on PDMS and cochlear implants

All zwitterionic thin film pre-polymer solutions were composed of 35wt% monomers, 0.8wt% surfactant, 0.05wt% Irgacure 2959 photoinitiator, and the remainder distilled water. The 35wt% monomer component was comprised of PEGDMA (crosslinker) and zwitterionic monomer (CBMA) to reach a desired percentage of crosslinker ranging from 0-100%. This prepolymer solution was used to coat medical grade PDMS in the form of sheets, small tubes, and cochlear implant devices for *in vivo* experiments. Initially, the PDMS pieces were placed in a 50 g/L benzophenone (surface grafting agent) acetone solution, soaked for 1 hour, and dried in vacuum.

Prepolymer solution was then dispersed over the surface using a cover slip for flat surfaces. For curved tube surfaces (1.2 mm outer diameter) an additional untreated PDMS sleeve (1.5 mm inner diameter) with a longitudinal slit was used to evenly disperse the solution. The photografted hydrogel thin film was then formed by photopolymerization using a UV lamp for 10 minutes at 30 mW/cm^2^ under nitrogen flow to mitigate any oxygen inhibition. Following polymerization, the cover slip or untreated PDMS outer tube was removed under wet (distilled water) conditions. The coated PDMS was stored in phosphate buffered saline (PBS).

The same coating method described above was applied to cochlear implants with minor modifications. Mid-scala and lateral wall cochlear implant electrode arrays were provided by Advanced Bionics (Santa Clarita, CA) and Cochlear (Sydney, Australia) respectively and were coated as above by insertion into a 0.76 mm inner diameter untreated PDMS tube with a longitudinal slit, filled with CBMA prepolymer solution incorporating 10wt% cross-linking monomer. To verify that the samples were successfully coated with the zwitterionic hydrogel, sodium fluorescein dyes of 20 mg/mL (concentrated) and 2mg/mL (dilute) were used.

### 2.3 Implantation surgery

Samples were implanted *in vivo* in 8-10 week old CBA/J mice (Fig. 4 – Fig. 6) or BL/6 mice (Fig. 3). Prior to any surgery, the implants were sterilized with UV light and surgical instruments were sterilized in an autoclave. All surgeries were conducted according to approved IACUC protocols, including proper anesthesia, analgesia, and post-surgical monitoring. After isoflurane anesthesia of 1-5% as needed by pedal response, an incision was made in the interscapular space and the samples were inserted under the dermis. Incisions were closed with absorbable sutures. The mice were monitored for post-surgical complications, and any mice developing ulcerative skin lesions were euthanized.

### 2.4 Scanning electron microscopy (SEM) imaging

After removal of the implants from mouse tissue, implants were soaked three times in PBS for 1 hour each. The samples were then cut in cross-section, dried in ambient air for 2 days, and mounted with the cutting face visible onto carbon tape. The sample was then gold sputtered and visualized under 10 kV of voltage at a working distance of 7.8 mm and an emission current of 93000 nA.

### 2.5 Confocal imaging of thin films

After removal of the implants from mouse tissue, the implants were soaked three times in fresh PBS for 1 hour each. The samples were then soaked in 20 mg/mL sodium fluorescein solution for 2 hours, removed, blotted dry, and placed on a microscope slide to image with a confocal microscope (Leica Stellaris 5 confocal with DMI 8). The settings for fluorescent imaging were calibrated using a positive control; a 4.5 μL drop of sodium fluorescein was dispersed under a 12 mm diameter glass coverslip to yield a fluid of 40 μm thick, and the gain adjusted so that the imaged confocal thickness matched the theoretical thickness.

### 2.6 *In vitro* fibroblast adhesion

Fibroblast adhesion on both naïve and explanted materials was determined using methods described previously.[24] Briefly, fibroblasts were harvested from p2-5 CBA/J mice and treated with 0.12% trypsin and 0.2% collagenase for 10 minutes at 37°C followed by treatment with fetal bovine serum (FBS) and Dulbecco’s Modified Eagle Medium (DMEM). Cells were then plated and cultured in DMEM 10% FBS. Confluent fibroblasts were removed from the cell culture with TrypLE Express. Samples were then exposed to 1 mL of resuspended fibroblasts for 48 hours at 37°C. Cultures were fixed with 4% paraformaldehyde in PBS followed by immunostaning with anti-vimentin antibody (Abcam ab92547, 1:200) to label the fibroblasts followed by a secondary antibody (ThermoFischer, 1:400) labeled with Alexa 488. The mounting medium contained DAPI to label nuclei. Fibroblast numbers were determined by capturing nine 20X images systematically per sample. For each sample, the mean number of fibroblasts per viewing field was averaged from the nine fields. Each condition was repeated with three samples.

### 2.7 Implant capsule tissue preservation for histology and immunohistochemistry

To harvest implants for histology and immunohistochemistry, the mice were perfused with 4% paraformaldehyde in PBS and treated with a depilatory agent. The implants were then excised together with the overlying skin and subdermal tissues. Samples were fixed with refrigerated 4% paraformaldehyde in PBS overnight, then washed three times for 1 hour each in PBS. The samples were then prepared for 1 hour each in successive PBS solutions of 10wt%, 20wt%, and 30wt% sucrose. Finally, the samples were placed in a 50:50 volume mixture of 30wt% sucrose in PBS embedding medium (Tissue-Tek OCT Compound) overnight at 4°C. The samples were then flash frozen in liquid nitrogen and sectioned on a LEICA CryoJane system to create 20μm thick sections. Sections were collected from a plane 3 mm deep from the leading edge of the implant to avoid edge effects.

### 2.8 Histology capsule thickness analysis

Sections stained with hematoxylin and eosin (H&E) were imaged with a light microscope at 4x and 20x magnification. The skin-facing side of the implant was systematically analyzed for capsule thickness at 20x magnification by measuring in ImageJ the distance from the implant interface through the basophilic cell layer to the transition from dark, thick eosinophilic (pink) banding to light eosinophilic banding.

### 2.9 Cochlear implant post-explant preparation, embedding, and sectioning

To prepare tissues encasing cochlear implant electrode arrays for sectioning, a modified epoxy embedding protocol was followed and is detailed in the Supplemental information. Sections were cut with a diamond knife at a thickness of 1 micrometer and stained with Toluidine Blue. Sections were collected from the cross-section of an exposed electrode interface. The presence or absence of grafted coating was confirmed by light microscopy of the sections to form groups of uncoated and pCBMA coated electrodes. Capsule thickness was assessed as described above, treating the regions of exposed PDMS and platinum separately.

### 2.10 Statistical methods

The confidence interval required for significance was set at 95% (p<0.05). In experiments with more than two independent variables, a two-way ANOVA was first conducted to detect which variables, if any, accounted for meaningful differences. If the threshold of statistical significance was met, then post-hoc Tukey tests were conducted to compare groups within statistically significant variables. One-way ANOVA was conducted on experiments with one independent variable and more than two groups, and follow-up Tukey tests conducted on statistically significant variables. For experiments comparing just two groups, Student’s t-test was used.

## Results

Surface modification is a promising avenue for reducing the foreign body response to biomedical implants while preserving inherent function. Such a modification needs to be durable, compatible with existing designs, and importantly, anti-fibrotic. To investigate the feasibility of pCBMA thin films as an anti-fibrotic surface on implants, systems were photopolymerized and photografted onto small PDMS implants and cochlear implant electrode arrays to assess *in vivo* durability and the impact of these coatings on fibrotic capsule thickness.

### 3.1 Characterization of pCBMA grafted thin films

Photografted zwitterionic pCBMA thin films coatings have previously been shown to photograft effectively to PDMS surfaces.[21] These coatings were applied to a variety of surface geometries and characterized as shown in Fig. 1. Fig. 1A demonstrates the hydrophilic properties of the zwitterionic coating, which readily absorbs the water solution colored with sodium fluorescein dye, as compared to uncoated PDMS, which fails to absorb the dye solution. To determine if these coatings could be applied to cochlear implant materials and persist *in vivo*, coated and uncoated PDMS sheets were placed subcutaneously in mice for 16 days. Samples were then removed, washed, and exposed to sodium fluorescein dye for confocal imaging. A hydrophilic zwitterionic coating should readily swell with the dye solution as compared to uncoated hydrophobic PDMS. The uncoated PDMS (Fig. 1C) shows essentially no absorption of dye, whereas fluorescence from the dye in the coating (Fig. 1D) shows maintenance of an approximately 40 μm thick and uniform film even after 16 days *in vivo*. For longer-term analysis of thin film persistence, coated and uncoated PDMS sheets were implanted subcutaneously for six months and then imaged by SEM, using cross-sections to evaluate for the presence of the thin film as shown in Fig. 1B-C. The thin film appears uniform and undamaged even after extended implantation suggesting that the hydrogels are durable and remain adhered to the PDMS *in vivo*. These same zwitterionic coatings were also photografted to lateral-wall cochlear implant electrode arrays. Images of uncoated (Fig. 1F) and coated (Fig. 1G) arrays are shown after being soaked in dye solution. A thin, uniform coating over the surface of the whole electrode array is evident, demonstrating the ability to coat cylindrical surfaces. Critically, the coating spans both regions of the PDMS housing and platinum electrodes.

**Fig. 1.**
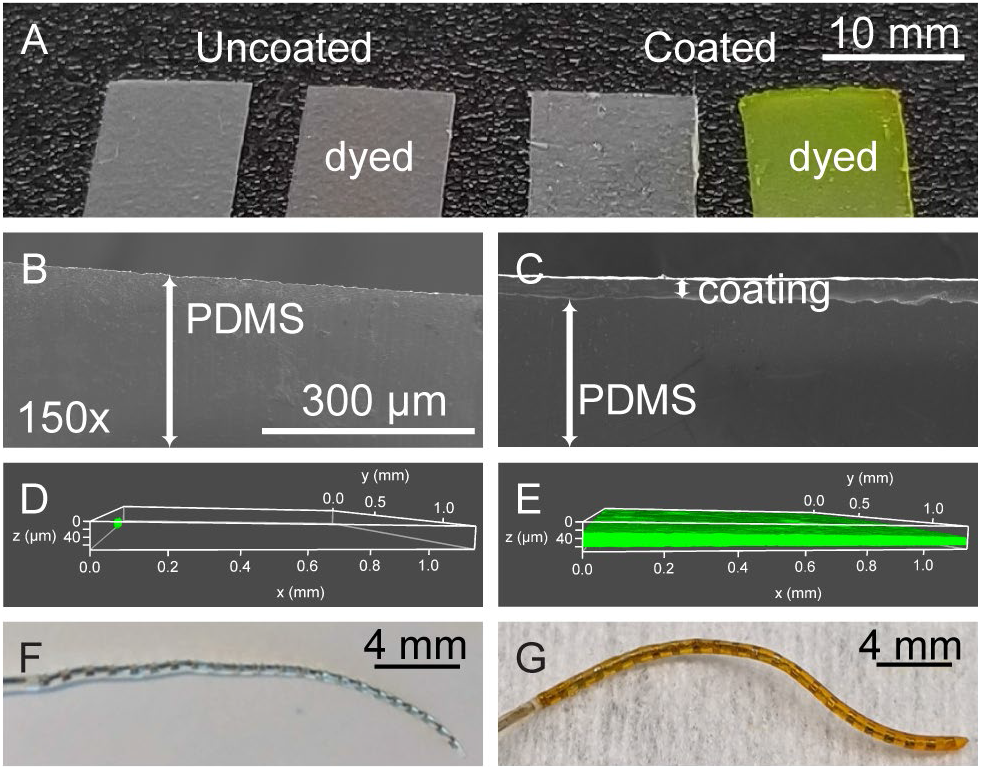
Photografted thin film zwitterionic hydrogels on PDMS. Shown from left to right are (A) images of uncoated PDMS, uncoated PDMS stained with fluorescein dye, coated PDMS, and coated PDMS stained with fluorescein dye (green). Additionally, SEM images of (B) uncoated PDMS and (C) coated PDMS both after 6-month implantation in subcutaneous mouse tissue in addition to confocal microscopy images of (D) uncoated PDMS and (E) coated PDMS both after 16-day implantation in subcutaneous mouse tissue and stained with fluorescein dye (green) are shown. Photographs of straight cochlear implant electrode arrays that are (F) uncoated and (G) coated and stained with concentrated fluorescein dye (orange) are also included. Scale bar in (B) applies to (C).

### 3.2 Fibroblast adhesion *in vitro* after subcutaneous incubation

To determine the effect of pCBMA cross-link density on anti-fouling properties, *in vitro* fibroblast adhesion was assessed on pCBMA coatings with 1.6%-31% cross-linker either freshly made or after six months of subcutaneous incubation. Shown in Fig. 2 are representative images of fibroblast adhesion to PDMS (Fig. 2A) and to PDMS coated with pCBMA hydrogel thin films of increasing crosslinker (Fig. 2B-E) with results quantified in Fig. 2F.

**Fig. 2.**
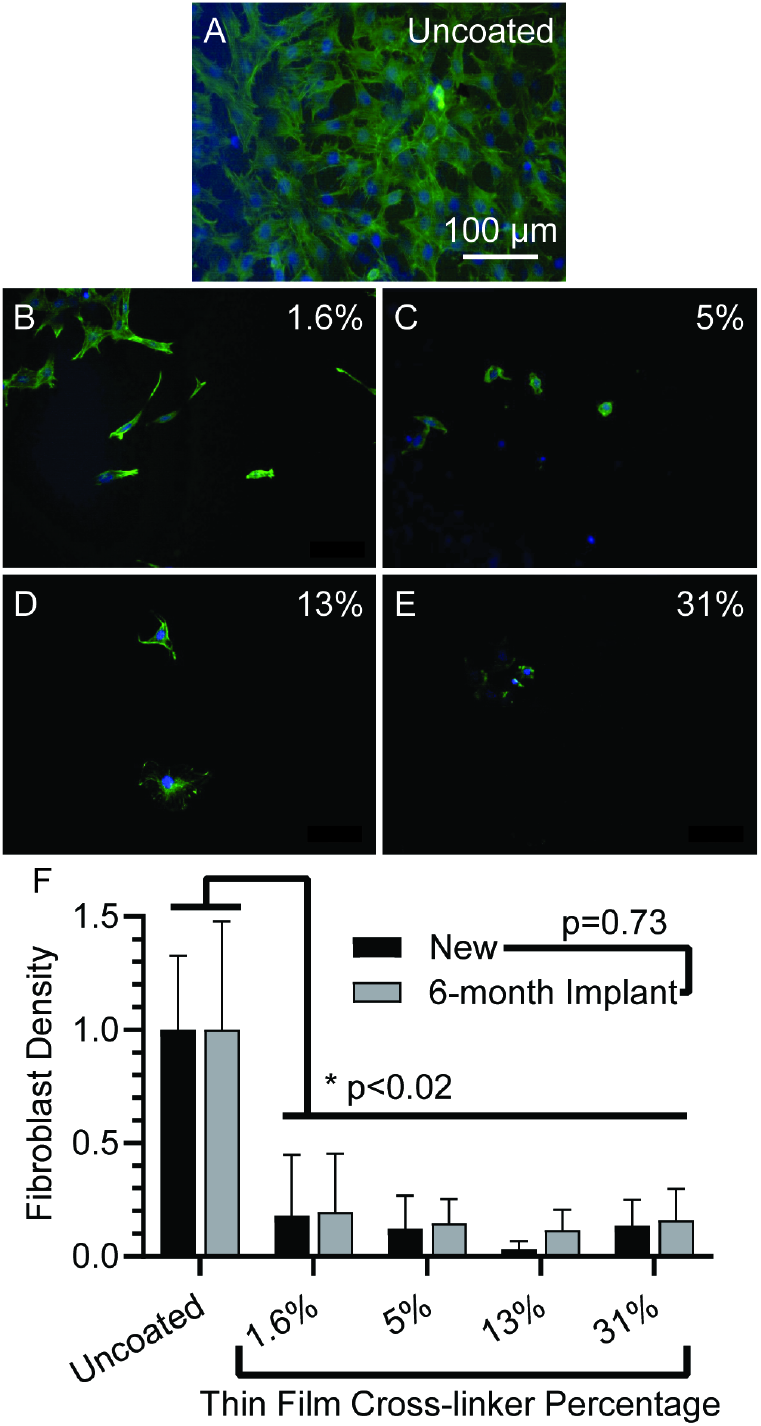
Photografted thin film zwitterionic hydrogels reduce in vitro fibroblast adhesion when initially prepared and after subcutaneous implantation for 6 months. Representative images are shown of fibroblasts labeled with antivimentin antibody (green, to stain fibroblast cell body) and DAPI (blue, to stain nuclei) as they adhere to (A) explanted uncoated PDMS and explanted PDMS that has been coated with pCBMA thin films of (B) 1.6%, (C) 5%, (D) 13%, and (E) 31% cross-linker percentages. (F) Fibroblast adhesion density is compared between uncoated (pair of leftmost columns) and pCBMA-coated PDMS of specified cross-linker percentages. Two-way ANOVA calculated a significant effect with surface modification. Each coated surface was compared against its respective uncoated control by Tukey’s post-hoc multiple comparisons test, and all are found to be significant (p<0.02). Also compared were new (black) and 6-month implants (grey) paired by surface type via two-way ANOVA. These two groups were not significantly different from each other (p=0.73). Error bars represent the standard error of the mean with n=3. Scale bar applies to all images.

The fibroblasts on the uncoated PDMS (Fig. 2A) appear as a thick, confluent mat of well- attached cells. The cellular processes are outstretched with relatively large cell bodies, rich in vimentin (green) staining. Very few fibroblasts are found on the coated surfaces (Fig. 2B-E). In addition to being much less dense, these cells display a more muted and compact morphology. These findings suggest that fibroblasts on the zwitterionic thin films lack the ability to readily attach and grow on the underlying surface. This effect is not from cytotoxicity, but because the cells do not adhere well.[25]

Accordingly, fibroblast adhesion to these coated surfaces is an order of magnitude less than the adhesion to PDMS (Fig. 2F). Two-way ANOVA revealed that the difference in cell adhesion between uncoated and cross-linked thin films was significant. Additionally, all the pCBMA systems with various degrees of cross-linking differed significantly from the uncoated control with a p value of 0.02 or less by post-hoc Tukey tests. This provides evidence that all pCBMA thin films, regardless of cross-linker percentage, demonstrate durable and effective anti-adhesion properties. The two-way ANOVA also showed no significant differences in fibroblast adhesion between new samples and explanted samples (p=0.73), confirming that these properties persist even after long-term subcutaneous implantation.

Altogether, these fibroblast anti-adhesion results provide evidence that photografted zwitterionic thin films retain their anti-fouling properties even after the mechanical, chemical, and biological insults of a long-term implantation and removal. Further, the hydrogel appears durable and effective against fibroblast adhesion over a wide range of cross-linker compositions.

### 3.3 Fibrotic capsule thickness and morphology

To determine whether macrophage [24] and fibroblast anti-adhesion properties translate to a reduced foreign body response *in vivo*, PDMS implants with both coated and uncoated regions were inserted into mice and removed after 6 weeks. The responses to uncoated regions differed substantially from the coated regions as shown in Fig. 3. The response to the uncoated region on the left of Fig. 3A is robust and highly cellular as depicted by a clearly defined dark basophilic (purple) interface next to thick, fibrous, highly cellular tissue. However, after progressing past the transition between coated and uncoated regions (shown in greater detail in Fig. 3B), the inflammatory response is only sustained for a few hundred micrometers and then dissipates. The interface with the pCBMA coating then continues without the intense cellular and fibrotic tissue found in the uncoated region. Thus, the pCBMA coating has a clear anti-fibrotic effect when directly compared in the same implant with a region of uncoated PDMS.

**Fig. 3.**
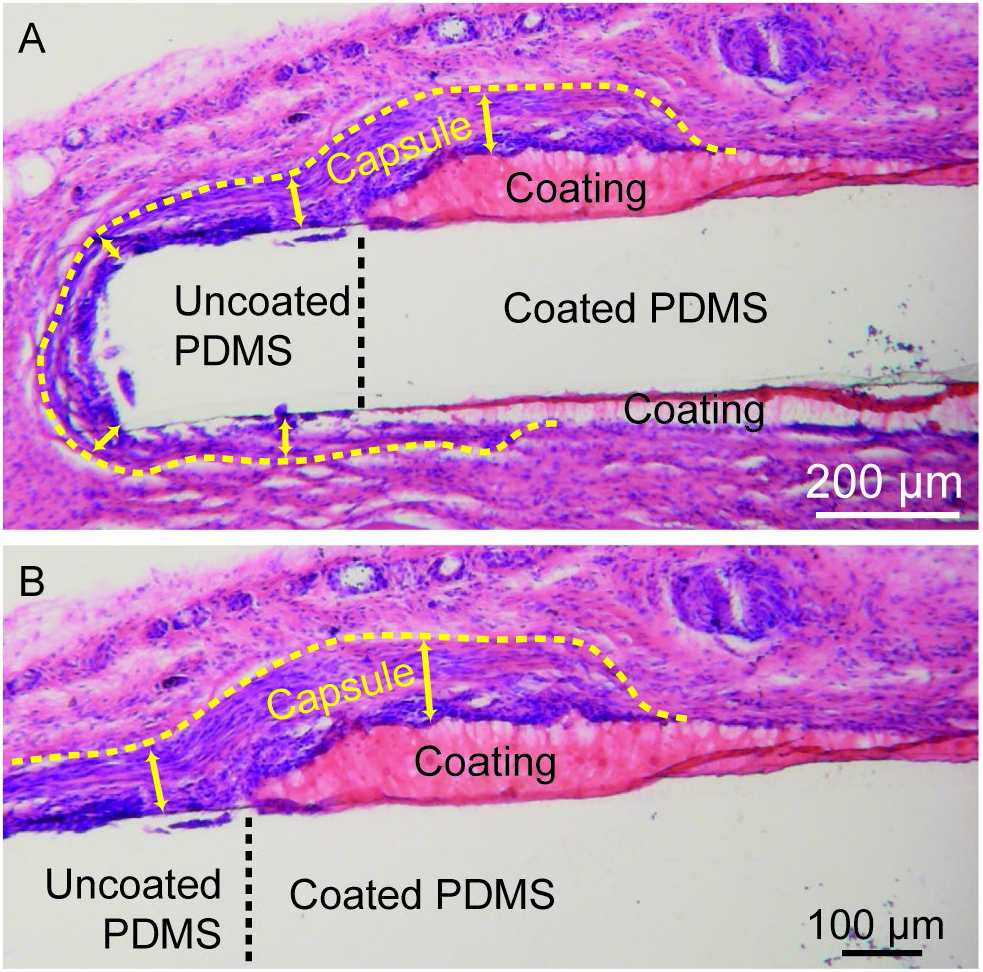
Capsule thickness is reduced, and the tissue morphology indicates less inflammatory response on pCBMA-coated regions of PDMS sheets. PDMS sheets with uncoated and pCBMA- coated regions were implanted in the subcutaneous tissue of mice. After 6 weeks the implants were harvested, and sections containing the implants were stained with hematoxylin and eosin. The pCBMA coating stains with eosin (pink) whereas the PDMS does not stain. Yellow dotted lines outline the fibrotic capsule, and the arrows indicate thickness. The vertical black dotted line indicates the boundary between the coated and uncoated region. (A) A wider and (B) more magnified view of the same image are included.

To quantitatively analyze the impact of photografted thin films on the fibrotic response, implants were inserted subcutaneously in mice. Fig. 4 shows the resulting histology and capsule thickness analyses. Implants tested included uncoated PDMS controls (Fig. 4A) in addition to poly[poly(ethylene glycol dimethacrylate)] films (pPEGDMA, Fig. 4B) and 10% cross-linked pCBMA films (Fig. 4C), all subcutaneously implanted for six weeks and histologically analyzed for capsule thickness (indicated with a dotted yellow line and asterisk in Fig. 4D). It should be noted that 10% cross-linked poly(hydroxyethyl methacrylate) (pHEMA) films were also prepared, but this mouse cohort developed ulcerative dermatitis surrounding the implant sites and required euthanasia within one week of implantation. This reaction to the implant, while lacking histology at six weeks, suggests that pHEMA thin films exacerbate the inflammatory response to the implanted materials rather than providing any degree of mitigation.

**Fig. 4.**
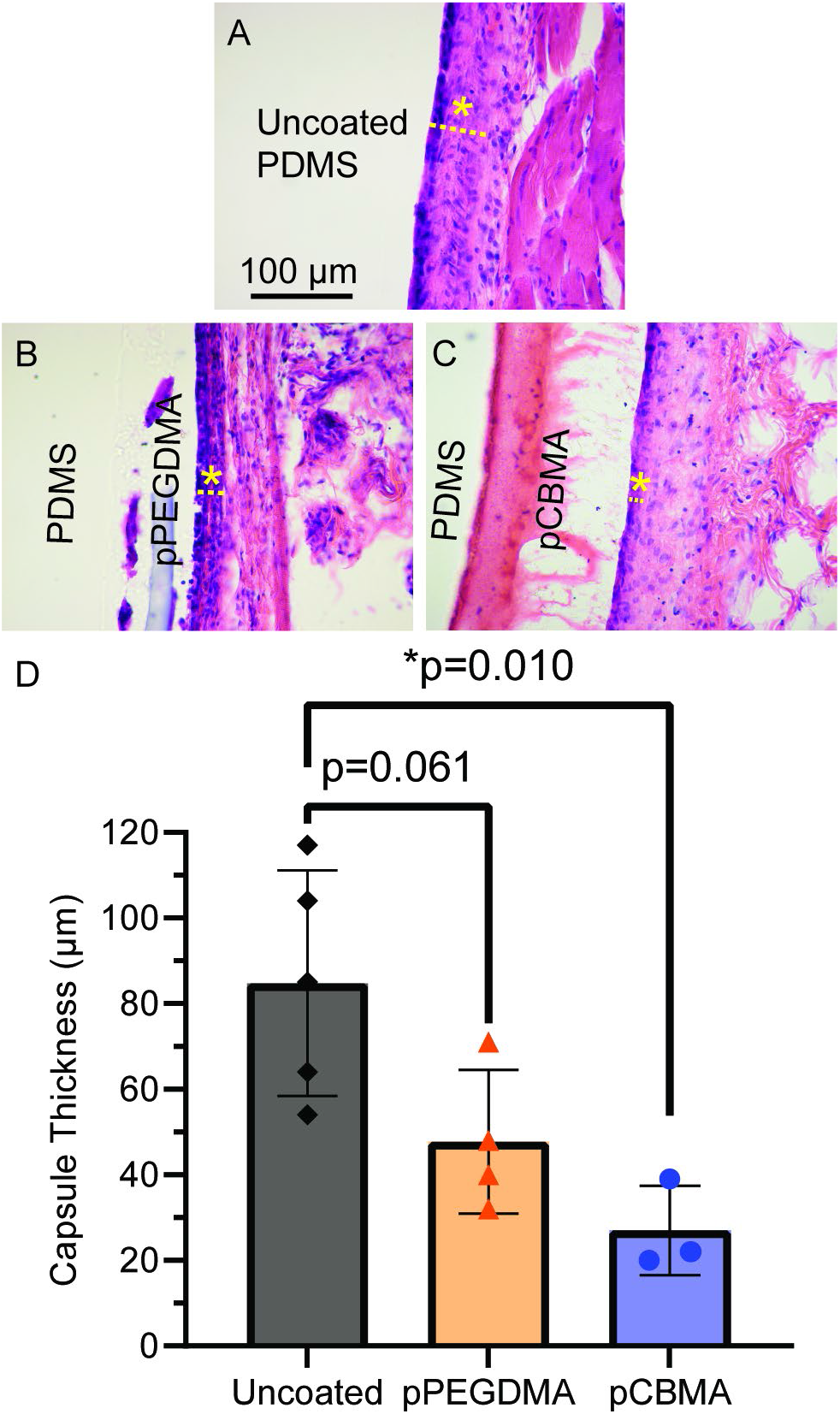
Photografted thin films of pCBMA more effectively reduce capsule thickness on uncoated PDMS sheets than pPEGDMA thin films. Images at 20x magnification are shown of the fibrotic capsule, with thicknesses indicated with a yellow dashed line and asterisk, after 6 weeks in vivo on PDMS sheets that were (A) uncoated, (B) coated with 100% pPEGDMA, and (C) coated with 10% cross-linked pCBMA. Tissue samples were stained by H&E to show basophilic cells (purple) and eosinophilic (pink) regions of fibrous tissue. One-way ANOVA revealed significant differences between the mean capsule thicknesses of uncoated versus pPEGDMA versus pCBMA (D). Post-hoc Tukey tests show that pCBMA and uncoated PDMS capsule thicknesses are significantly different (p=0.010) while pPEGDMA does not show any statistical difference to uncoated PDMS (p=0.061). Error bars represent the standard error of the mean with n=3-5. Scale bar applies to all images.

The capsule on the pCBMA coated implant shows much less foreign body response pathology than either of the controls. The accompanying capsule is much less pronounced, with a mean thickness of 27 μm compared to 82 μm for uncoated and 43 μm for pPEGDMA (Fig. 4D). Statistically, capsule thickness is significantly different when comparing uncoated, pPEGDMA, and pCBMA (Fig. 4D) by one-way ANOVA. Additionally, post-hoc Tukey tests reveal that the capsule thickness decrease in pCBMA relative to uncoated PDMS is significant (p=0.010), and that the pPEGDMA narrowly fails to reach the threshold of significance (p=0.061).

On uncoated PDMS (Fig. 4A), a dense infiltrate of basophilic cells immediately at the interface is evident. These cells dominate the otherwise lightly eosinophilic (pink) and fibrous connective tissue with densely packed basophilic bodies. Directly adjacent to this basophilic milieu, a band of intense eosinophilic stain reveals tightly packed fibers running parallel to the interface. This fibrous tissue forms a continuous barrier around the implant. Moving away from the dense fibrous tissue towards the skin, the tissue suddenly stains less eosinophilic, fewer cells are present, and connective fibers appear loosely organized and randomly oriented, which is typical of loose connective tissue found in normal subdermal tissue. The thickness measurements and analysis therefore focus on the pathology of the inflammatory, dense, fibrous capsule.

When the implant is coated with a thin film of zwitterionic hydrogel (Fig. 4C), the appearance of the implant-tissue interface is quite different. First, the hydrogel itself is clearly visible as a bright, uniform, acellular, eosinophilic band under H&E staining. The interface with the tissue begins at the edge of this band where cells first appear. At this interface, the tissue either appears as loose connective tissue or as a one or two nuclei-wide basophilic band which quickly gives way to a narrow band of deeply eosinophilic fibers outwards toward the skin. Typically, this band is limited in scope or even absent. Overall, the tissue response immediately around the zwitterionic coated system appears much less basophilic and cellular than the tissue seen around an uncoated implant. The adjacent tissue moving away from the thin film stains much more lightly eosinophilic. Taken together, these findings indicate a thinner capsule and diminished foreign body response to the pCBMA coating.

The grafted thin film composed of cross-linker molecules only (pPEGDMA) was tested as a non-zwitterionic hydrogel control. The scar capsule and morphology are interestingly more like those found in uncoated PDMS rather than those seen in response to the pCBMA thin film. pPEGDMA was selected as a control both because it is used in small amounts (10-20%) as a cross-linker in the pCBMA films and also a well-studied biomaterial.[26] The immediate interface appears as an intense, thick, basophilic region with multinucleated giant cells (Fig. 4B). The hydrogel itself is visible as a relatively thin, cloudy, blue-gray band that appears to be missing in regions across the surface, suggesting that the pPEGDMA hydrogel likely does not adhere effectively in all regions to the underlying PDMS. Where the film is present, a thin line of cells is observed between the PDMS and pPEGDMA film, implying that adhesive failure of the film likely occurred during the implantation period and that the void formed *in vivo*. To this point, all films were confirmed to be intact and defect-free prior to insertion. Therefore, the pPEGDMA film appears unable to fully withstand the *in vivo* environment. Altogether, these data show that pCBMA coatings more robustly mitigate the foreign body response on PDMS than coatings made primarily from other conventional biomaterials like PEGDMA and HEMA.

### 3.4 Fibrotic capsule thickness as a function of cross-link density

Previous work showed that the *in vitro* anti-fouling properties were a strong function of cross-link density with the greatest enhancement between 5-50% cross-linker.[24] To determine if capsule thickness and morphology are also a function of cross-link density, pCBMA thin films of 0-50% cross-linker were subcutaneously implanted for six weeks and analyzed by H&E staining. Additionally, it is also important to assess whether the photografting process and coating durability could be effective *in vivo* with cylindrical substrate geometry. Following the implantation, capsule thickness was analyzed as shown in Fig. 5. The uncoated control (Fig. 5A) histology reveals a well demarcated band of fibrosis encompassing the implant (marked with a dotted line and asterisk) including a layer of dense, basophilic cells immediately at the interface, consistent with previous results. Some of the nuclei at the surface appear conglomerated into giant cells, a typical development of the late-stage foreign body response.

**Fig. 5.**
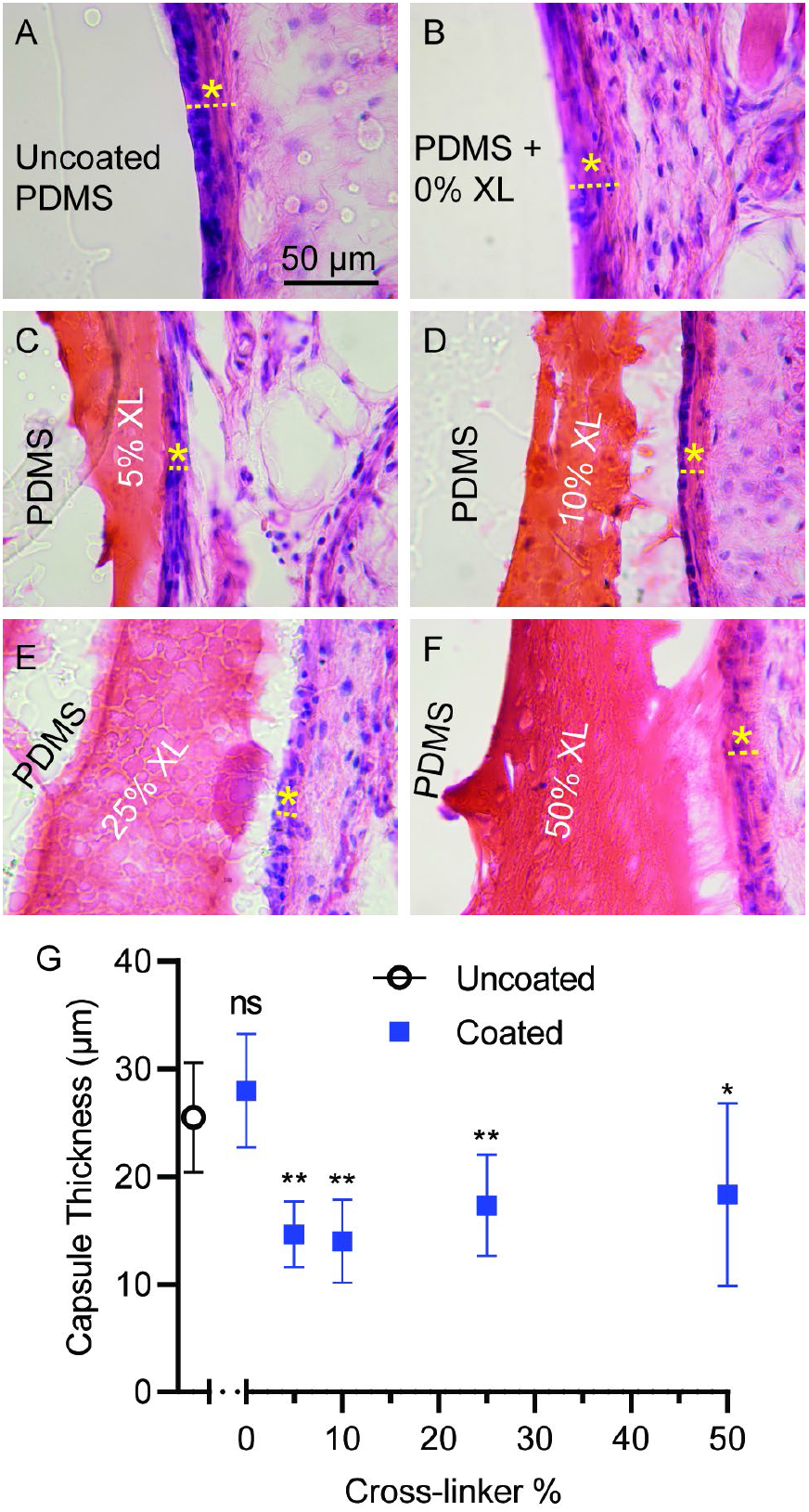
Photografted pCBMA hydrogel thin films with a broad range of cross-linker percentages reduce fibrotic capsule thickness on cylindrical PDMS. Images at 20x magnification are shown of fibrotic capsule thicknesses (labeled with a yellow dashed line and asterisk) after 6 weeks in vivo on PDMS cylinders of (A) uncoated PDMS and pCBMA coated samples of (B) 0%, (C) 5%, (D) 10%, (E) 25%, and (F) 50% cross-linker. Tissue samples were stained by H&E to show basophilic cells (purple) and eosinophilic (pink) regions of fibrous tissue. (D) The graph plots the mean capsule thickness as a function of surface type, with significance of a comparison to PDMS summarized “ns” for not significant, “*” for p<0.05, and “**” for p<0.01. One-way ANOVA yielded a significant difference between groups at p=0.0044 and was followed by Dunnett’s multiple comparison test, using uncoated PDMS as the control. Compared to the uncoated system, the cross-linker percentage is not significant for 0% (p=0.3322) but was significant for 5% (p=0.0039), 10% (p=0.0031), 25% (p=0.0031), and 50% (p=0.0265). Error bars represent the standard error of the mean with n=3. Scale bar applies to all images, all imaged at 20x magnification.

The deep, intense eosinophilic staining of the overall capsule sharply contrasts with the lightly stained surrounding tissue.

A control without cross-linker was also examined, because this formulation is expected to only form relatively weak pCBMA strands grafted from the surface. Such systems have shown greater fouling *in vitro* compared to robust pCBMA cross-linked hydrogels.[24] As expected, this control lacked evidence of any significant coating after sectioning (Fig. 5B) and elicited a robust capsule of similar thickness to uncoated PDMS. The capsule itself matches the intense eosinophilic banding seen in the uncoated sample, although without the same degree of interfacial basophilic cells. This result supports our previous *in vitro* findings that cross-linker is necessary to enable signficant anti-fouling properties on a surface via dense, networked, grafted pCBMA polymer chains.[24]

The capsule is quite different, however, at the interface with cross-linked pCBMA thin films (Fig. 5C-F). Starting with the pCBMA thin film of 5% cross-linker (Fig. 5C), the thin film itself stains an acellular orange and interfaces with a limited band of cellular, basophilic fibrosis. Interestingly, the coloring and stain of these cells lack the intensity seen in the uncoated sample. The capsule thickness is also reduced by a factor of 2 compared to both uncoated and uncross-linked samples. Similar observations apply to the pCBMA coating of 10% cross-linker (Fig. 5D), although this capsule appears to have far fewer interfacial cells and is mostly a thin band of eosinophilic capsule. The 25% cross-linker capsule (Fig. 5E) is less defined. The material interface exhibits a faint band of increased eosinophilia and slightly more concentrated cells, but appears the most similar overall to normal loose connective tissue. As the film becomes more cross-linked and richer in PEGDMA, it is reasonable to believe that characteristics will more closely match that of the pure PEGDMA film from Fig. 4B. We find that the 50% cross-linked CBMA film (Fig. 5F) remains intact, shows bright staining on H&E, and mitigates capsule development, although not as consistently or strongly as lower cross-link percentages. The capsule bears a slight enrichment in cells and demonstrates eosinophilic banding, but not as intensely as the uncoated sample.

One-way ANOVA indicated a statistically significant variation between groups (p=0.0044). Follow-up Dunnett’s multiple comparison tests compared each group to the uncoated control. While the 0% cross-linker implant capsule appeared the same statistically as the one on uncoated PDMS (p=0.33), all cross-linked films had a statistically significant decrease in their mean capsule thickness compared to uncoated.

These results together suggest that a wide range (5%-50%) of cross-linker percentages can be used in pCBMA thin films to reduce the foreign body response on surfaces, including those from cylindrical geometries. The reduction is seen most effectively and consistently between 5%-25% cross-linker. Cross-linking in this system is necessary to form a hydrogel layer that comprehensively protects a surface *in vivo*, since pCBMA photografted with no cross-linker showed the same *in vivo* capsule thickness as an uncoated surface.

### 3.6 One-year fibrotic capsule thickness on human cochlear implant electrode arrays

To assess the *in vivo* anti-fibrotic effect of zwitterionic thin films on medical implants, pCBMA coatings were applied to cochlear implant electrode arrays and implanted subcutaneously for one year. Coating cochlear implant electrode arrays and assessing the capsule thickness formed *in vivo* represents a technical challenge. First, properly and permanently coating the exposed platinum electrode is difficult because typical metal grafting approaches rely on chemical modification at the surface, but the platinum electrodes are inert. The approach here leveraged the internal cohesion of hydrogel to span the platinum electrode regions from surrounding anchored hydrogel regions on PDMS. The second challenge is in preparing the tissues containing cochlear implant electrode arrays for sectioning. A specimen with an electrode array in fibrotic subcutaneous tissue demonstrates very different stiffnesses that make traditional sectioning techniques infeasible. The approach here used a stiff epoxy resin to embed the specimen followed by sectioning with a diamond knife to cut through the stiff, hard cochlear implant electrodes while still preserving the integrity of the surrounding tissue and implant housing.

Pre-curved mid-scala cochlear implant arrays were straightened with a stylet and photografted with pCBMA hydrogels. The stylet was then removed and, importantly, the thin film withstood the recurving of the electrode array. This is significant since electrode arrays adopt a curved configuration when placed within the spiral shaped cochlea. Coated and uncoated mid-scala cochlear implant arrays were then implanted subcutaneously for one year to determine the degree of foreign body response in a long-term setting. Over this time frame, any foreign body response should reflect the lifetime impact. The implants were then removed, prepared for embedding, photographed (Fig. 6A-B), and embedded in resin (Fig. 6C-D) for sectioning.

**Fig. 6.**
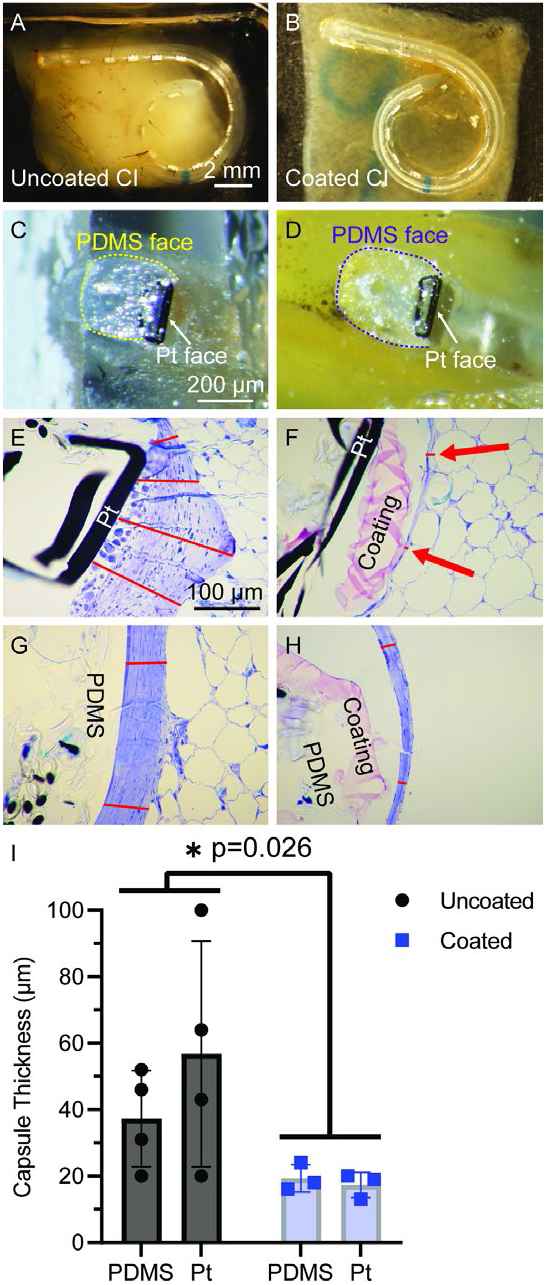
Photografted pCBMA thin films reduce 1-year in vivo fibrotic capsule thickness subcutaneously on human cochlear implant electrode arrays. The left column (A, C, E, G, and gray bars of I) show images and results from uncoated electrodes and the right column (B, D, F, H, and blue bars of I) of pCBMA-coated electrodes. The top row shows images of cleared (A) uncoated and (B) coated cochlear implants after 1 year of implantation in subcutaneous mouse tissue and shortly before embedding in resin. The second row (C and D) shows images of (C) uncoated and (D) coated cochlear implant electrode arrays embedded in resin and cut in cross-section in the middle of an exposed platinum electrode (labeled “Pt face”) and PDMS housing (labeled “PDMS face”). The third row (E and F) shows 20x magnification images of 1 μm sections of (E) uncoated and (F) coated (coating is pink in appearance and labeled “Thin Film”) platinum electrode faces (labeled “Pt”) stained with Toluidine blue, with the fibrotic capsule thickness indicated by red lines. In the coated electrode, the thickness is much smaller, so its capsule is indicated with red arrows. The 4th row shows similar 20x magnification images, but of the PDMS-housing face of the implant, again stained with Toluidine blue and the capsule thickness indicated in red for (G) uncoated and (H) coated (coating which is pink in appearance is labeled “Thin Film”). (I) Capsule thickness is plotted as a function surface type (PDMS versus Pt, platinum-iridium) and surface treatment (uncoated shown as black circles versus coated shown as blue squares). Two-way ANOVA indicates a significant difference for uncoated vs coated groups of p=0.026. Error bars represent the standard error of the mean, n=3-4. Scale bars apply to subsequent images until a new scale bar is given.

Cross-sections were cut from electrode-bearing regions and were stained with Toluidine Blue. Sample images are shown in Fig. 6 for uncoated platinum electrodes (Fig. 6E), coated platinum electrodes (Fig. 6F), uncoated PDMS housing (Fig. 6G), and coated PDMS housing (Fig. 6H). The capsule thicknesses were measured for each electrode and were analyzed by two-way ANOVA (Fig. 6I). The capsule surrounding the uncoated platinum electrode is thick and appears almost pyramidal in shape, the apex being positioned over the middle of the electrode face. This effect suggests that immune interactions at the platinum interface produce a local signal for intense foreign body response activation that is maximized in the center of the exposed platinum.[17] The cellular architecture immediately at the interface of the platinum also suggests a sustained presence of immune cells. The fibrotic capsule is thick, banded, and clearly identified from the surrounding minimally stained adipose tissue.

The zwitterionic coating on the platinum (Fig. 6F) is easily identified by the presence of the pink-stained coating which is positioned between the black, opaque platinum face and the surrounding tissue. A minimal band of fibrosis is present, but the intense fibrosis observed in the uncoated platinum is largely absent. The thin film itself remains intact and spans the region over the platinum electrode, showing a uniform film that completely encapsulates both the PDMS and platinum-iridium surfaces. It should be noted that the platinum appears fractured as an artifact of sectioning stresses but remained intact during implantation as evidenced by the intact electrodes on the cutting faces of Fig. 6C-D.

A clear difference is also observed between the capsule on uncoated and coated PDMS housing. The fibrotic capsule of uncoated PDMS stains deep blue and exhibits the typical parallel banding of fibrotic capsules. The pCBMA coated system induces a much thinner capsule, as shown in Fig. 6H. It should be noted that due to the difficulty of sectioning, implant elements may have shifted slightly during sectioning, especially in terms of the exact positioning of the thin film relative to the tissue. The PDMS housing itself does not absorb any stain and so appears colorless.

Two-way ANOVA revealed a significant difference between coated and uncoated mean capsule thickness. The mean capsule thickness for an uncoated cochlear implant was 47 μm but was 60% less at 18 μm for a coated cochlear implant. While a statistical difference was not observed between PDMS and platinum surfaces, the platinum surface appears to have elicited a stronger response, consistent with previous studies.[17] Interestingly, the coating seems to mask the differences in the underlying surface material, and no difference was observed between coated platinum and PDMS surfaces. Very little variability was observed in capsule thickness for coated samples. On the other hand, variability was high in uncoated samples. With a higher-powered study, any interaction between coating status and difference between platinum and PDMS effects might become discernable. However, the goal of this experiment was to determine whether coating a cochlear implant electrode array showed a demonstrable effect on capsule thickness over a chronic implantation. In that respect, the study accomplished its aims.

We see noteworthy evidence that pCBMA coatings bridge platinum electrodes, maintain long-term durability, and effectively mitigate the foreign body response. The effect substantially reduces the chronic immune responses out to one year. It is reasonable to believe that this reduced fibrotic capsule would lead to higher signal resolution, lower impedances, and improved implant longevity for cochlear implants and other electrode-bearing implants.

## Discussion

This work demonstrates for the first time the *in vivo* anti-fibrotic effect of a photografted zwitterionic hydrogel thin film on medical device materials. Specifically, the capsule thickness surrounding coated materials is reduced and histologically shows fewer interfacial cells and reduced fibrotic tissue (Fig. 3 - Fig. 6). When conventional (pPEGDMA, pHEMA) and zwitterionic hydrogel thin films (pCBMA) are compared (Fig. 4), zwitterionic films exhibit a greater degree of durability, compatibility, and anti-fibrotic effects. In particular, pPEGDMA thin films do not remain intact over a six-week incubation *in vivo* and pHEMA films induce ulceration, whereas pCBMA films remain intact while reducing capsule thickness.

This work also highlights the ability to photograft zwitterionic thin films on flat (Fig. 1A, Fig. 3, Fig. 4) and cylindrical (Fig. 1G, Fig. 5, Fig. 6) geometries while maintaining anti-fibrotic properties. Multiple experiments show that the anti-fouling effect of zwitterionic hydrogels is largely independent of cross-link density (Fig. 2, Fig. 5) as long as zwitterionic monomers account for at least half of the composition of the polymeric hydrogel. These results also demonstrate that thin films are both physically present and functionally unchanged *in vivo* at durations of six months (Fig. 1, Fig. 2) and one year (Fig. 6).

The results suggest two potential independent mechanisms for the change in tissue response to a coated surface. The mechanism best supported by the data is that decreased attachment of fibroblasts (Fig. 2) and other immune cells drives the reduction in fibrosis seen on coated implants. Decreased adhesion of any agent of the foreign body response, whether fibrinogen, macrophage, or fibroblast, would be expected to dampen the inflammatory cascade necessary for producing a fibrotic capsule.

While the anti-adhesion properties of zwitterionic thin films are likely contributing significantly to this capsule-reducing effect, another potential mechanism could also be active. The microenvironment at the interface of a coated implant may mimic the endogenous environment better than that of an uncoated implant. If this microenvironment better resembles native tissues, it would likely trigger less inflammatory interactions and local signaling. Potential evidence of this effect can be seen in Fig. 3, where regions of coating that closely border uncoated regions still receive local inflammatory signals, while those regions far (e.g. more than ~600 μm) from uncoated sites appear relatively protected from pro-inflammatory signaling. This mechanism could also account for reduced capsule thickness even when cells do ultimately colonize the implant interface.

Despite the historically difficult nature of permanently modifying inert metal surfaces, this coating innovation demonstrates the ability to mitigate capsule formation around platinum electrodes (Fig. 6). This result is promising not just for cochlear implant electrode arrays, but also for any electrode-based device that would benefit from an *in vivo* anti-fibrotic effect such as glucose monitors and deep brain stimulators. In this work, the thin films benefitted from anchoring regions of PDMS housing to help adhere to these electrodes and proved effective for covering ~0.5 mm electrode leaflets that were recessed into the PDMS housing. The evidence suggests that the film remained flush with the electrode surface, otherwise it would have seen cellular infiltration in the gap like was observed with imperfectly adhered pPEGDMA films (Fig. 3B).

For cochlear implants in particular, a capsule thickness reduction of 50-70% could have dramatic implications for auditory performance. Reducing fibrosis could help mitigate the gradual increase in electrode impedances that occurs in the months following implantation and lead to reduced current spread and improved battery life. Further, delayed loss of residual acoustic hearing after cochlear implantation has been associated with intracochlear fibrosis and increased electrode impedance.[27–30] Thus, to the extent that zwitterionic thin film coatings reduce fibrosis, they may also be expected to mitigate loss of residual hearing after cochlear implantation.[31, 32]

This work is based on a mouse subcutaneous model for assessing the extent of the foreign body response to implants, which offers some advantages and disadvantages. The subcutaneous model enabled an assessment of the foreign body response in isolation from the potentially confounding effect of cochlear trauma. Importantly, we have already shown that these zwitterionic thin films dramatically reduce surface friction of PDMS leading to a reduction in insertion forces for cochlear implant electrode arrays in human cadaveric cochleae.[22] A subcutaneous mouse model enables evaluation of anti-fibrotic performance of a human cochlear implant electrode array. Implants sized for humans were desirable because they were large enough to feasibly coat using the sleeve polymerization described in the methods. Human implants of course have an advantage over miniaturized implants because they best reflect actual clinical use. Further, they are available in pre-curved configurations to mimic the shape of the electrode array when placed in the cochlea. However, some elements of the subcutaneous model limit the study. For one, the impact of zwitterionic hydrogels on mitigating cochlea-specific responses like neo-ossification remains unknown. The biology or anatomy of the cochlea could possibly limit the functionality of a zwitterionic hydrogel in ways yet to be understood. Further, improvements in cochlear implant electrode array performance can only be explicitly assessed with a functional implant in a living cochlea. To address these limitations, future work will investigate the effect of zwitterionic thin films on fibrosis, electrical impedance, and neo-ossification using functional human cochlear implants implanted in cochleae of large animals.

## Conclusions

This study shows the potent anti-fibrotic effect of photografted pCBMA thin films on implant materials including PDMS in different geometries and cochlear implant electrode arrays. This study provides *in vivo* evidence that a wide range of cross-linker compositions can be effectively used in coatings to mitigate the foreign body response. These pCBMA coatings are durable and maintain their properties over six-month and one-year long incubations. The reduction in fibrotic capsule thickness offered by this passive, durable thin film is sustained *in vivo* up to one year, which is promising for implants needing long-term protection. For a cochlear implant, a coated electrode may lead to enhanced hearing performance and reduce risk of loss of residual hearing. These findings stand to benefit not just outcomes for cochlear implants, but also many other neural prostheses or other medical devices that would show enhanced performance or fewer complications with reduced fibrosis.

## Acknowledgements

This project was supported by the National Institutes of Health (5T32DC000040, T32GM139776, F30DC019274, and R01DC012578) and by the University of Iowa UI Ventures fund.

The authors would also like to thank Advanced Bionics for donating inactive mid-scala human cochlear implant electrode arrays and Cochlear for donating inactive Nucleus Slim Straight human cochlear implant electrode arrays. Additionally, the authors acknowledge Jianqiang Shao for extensive assistance with the sectioning and embedding of the explanted cochlear implant electrode arrays and Brian Mostaert for troubleshooting and designing effective tissue sectioning methods.

## Supplemental information

Samples were soaked in ½ strength Karnovsky’s fixative overnight at 4°C, treated with a 0.1M cacodylate buffer three times for 30 minutes each, transferred to a 1% OsO4 solution in 0.1M cacodylate buffer for 1-2 hours, transferred to a 1% OsO4 and 1.5% potassium ferrocyanide in cacodylate buffer solution for 1-2 hours, placed in 0.1M cacodylate buffer again 3 times for 20 minutes each, followed by rinsing with distilled water for one minute. The explants were then placed in 2.5% uranyl acetate for 20 minutes. Next, the tissues were soaked in a series of ethanol/water solutions of 50%, 75%, 95%, and 100% for 15 min, 15-20 min, 30 min, and 2 times for 30 min each, respectively. Next, the tissues were soaked in a 1:1 ethanol:propylene oxide (PO) solution for 30 minutes then pure PO for another 30 minutes. The explants were placed in a 2 parts PO and 1 part LRW epoxy mixture for 30 minutes then a 1 part PO and 2 parts LRW epoxy mixture for 1 hour. Then the samples were soaked in pure LRW epoxy twice for 30 minutes each. The cochlear implants were then embedded in fresh LRW epoxy that was prepared in flat molds and baked in a 70°C oven for 2 days.

